# A Low-Cost Mass Spectrometry-Based Approach for Quantifying Purines in Placental Extracts

**DOI:** 10.1101/2020.11.19.389817

**Authors:** Ruslan Rodriguez, Igor Konovets, Serhii Ralchenko, Maxsim Kharkhota, Andrij Kostyuk, Victoriia Kosach, Irina Voronina, Natalia Filimonova, Maria Obolenskaya

## Abstract

Hyperhomocysteinemia is a medical condition characterized by an abnormally high level of homocysteine in the blood associated with multiple human pathologies including preeclampsia – the most feared complication of pregnancy, with placenta playing the central role in the pathogenesis of preeclampsia. The developing placenta is highly sensitive to different adverse factors but its response to hyperhomocysteinemia is not fully clear. Previously we have demonstrated the activation of reactions of methionine cycle and the transsulfuration pathway in placental explants cultivated with homocysteine. The reactions of the methionine cycle are tightly connected with reactions of the folate cycle, encompassing reactions of *de novo* purine biosynthesis, which are crucial for the developing placenta, as they support rapid ATP generation to maintain energy status and increased biosynthesis of macromolecules. The sensitivity of *de novo* purine biosynthesis to hyperhomocysteinemia is not known. The aim of this study was to evaluate the impact of homocysteine on placental *de novo* purine biosynthesis.

**Methods:** We developed a simplified method to measure the level of all and newly formed purines by HPLC/ESI-MS, using a stable isotope glycine to label newly synthesized purines. The developed method proved to be highly sensitive, interday repeatable and intraday reproducible. We applied a method for placental explants from the first and third trimesters of gestation and MCF7 cells cultivated with 20 μM and 40 μM homocysteine corresponding to its concentrations at mild and intermediate hyperhomocysteinemia.

**Results:** The developed method proved to be highly sensitive, interday repeatable and intraday reproducible. The content of total purines in placental explants from the first trimester of gestation was around 9.0 μmol/g wet tissues at specified conditions of cultivation. The newly formed purines comprised around 1 % of total purines, decreased steadily in explants cultivated with 20 μM and 40 μM homocysteine, and reached the values characteristic for explants from third trimester cultivated without homocysteine, 4.0 μmol/g wet tissues. The effect of homocysteine reproduced with MCF7 cells.

**Conclusion:** Homocysteine in concentrations characteristic of mild and intermediate hyperhomocysteinemia induces the down regulation of *de novo* purine biosynthesis in placental explants, and implies the shift of metabolic pathway to homocysteine remethylation and transsulfuration at the expense of *de novo* purine biosynthesis.

## Introduction

Purine nucleotides and related phosphorylated metabolites have many important roles in the core processes of cellular metabolism. They are precursors in the synthesis of nucleic acids; ATP and GTP are energy carriers; derivatives of purine nucleotides are coenzymes (coenzyme A, NAD, NADP, and FAD) and regulators in cellular metabolism (cAMP and cGMP); they serve as signaling molecules operating through purinergic receptors. Purine nucleotides take part in metabolic regulation, as in the response of key enzymes of intermediary metabolism to the relative concentrations of AMP, ADP, and ATP [1, 2]. Purine biosynthetic activity modulates immune reactions and indirectly controls self-tolerance [3, 4].

Coordinated action of salvage and *de novo* biosynthetic pathways maintains purine levels in mammalian cells. Under normal physiological conditions, when most adult cells are at the resting state, the low-cost recycling of intermediate products of purine nucleotides catabolism via the salvage pathway maintains most of the cellular purine nucleotide pool. In contrast, during active proliferation and growth requiring higher purine levels, the up regulated *de novo* biosynthetic pathway and assembling of its six enzymes in multi-component complex purinosome satisfies intracellular purine demand [5]. The *de novo* pathway is a highly conserved and energy-intensive process. It uses five molecules of ATP, two molecules each of glutamine and 10-formyltetrahydrofolate (THF), and one molecule each of glycine, aspartate, and carbon dioxide for one molecule of inosine monophosphate (IMP), a common intermediate and branch point to ATP and GTP production [1]. Glutamine and aspartate are generated from intermediates of the tricarboxylic acid cycle; glycine and formate are derived from tetrahydrofolate (THF) -dependent serine and glycine catabolism in mitochondria [2, 6, 7]. Thus, *de novo* purine synthesis is connected with the core pathways of cellular metabolism; purine nucleotide pools are in bidirectional regulatory relationship with the activity of the mechanistic target of rapamycin (mTOR) complex, which functions as a nutrient/energy/redox sensor [8, 9].

Despite recent breakthroughs in the study of *de novo* purine synthesis, there are many processes which still require explanation: for example, the mechanisms of tissue and cell specific regulation of the synthesis of six enzymes, carrying out *de novo* purine biosynthesis, and the molecular mechanisms of response to extracellular agents. Homocysteine is an adverse extracellular agent, whose elevated concentration in blood (hyperhomocysteinemia) is associated with multiple human pathologies, and characterized by a tissue-specific effect on *de novo* biosynthesis [10, 11]. The *de novo* purine pathway is a promising target for anticancer therapy requiring detailed study.

At present, there are different approaches to assay *de novo* purine nucleotide synthesis, from quantification of newly synthesized nucleotides including acid soluble nucleotides and those incorporated in DNA and RNA nucleotides [12], to the comprehensive quantitative profiling of nucleotides and related phosphate-containing metabolites [13]. The protocols for estimation of *de novo* purine biosynthesis vary in the following ways: usage of various precursors: glycine, serine, formate or glucose; radioactive or non-radioactive (stable) isotopes for labeling; systems for labeled product detection: liquid scintillation counting, high performance liquid chromatography (HPLC), mass-spectrometry (MS or MS/MS), or nuclear magnetic resonance; and, finally, in the financial cost and time required. Although many methods are used to measure nucleotide levels, only a few reports are devoted to the quantitative tissue-specific characteristics of purine production, especially in low mg weight biological samples.

The present study aimed to develop and validate a simplified and low-cost method for purine quantification and estimation of *de novo* purine synthesis in human placenta in quasiphysiological conditions and in simulated hyperhomocysteinemia. HPLC with electrospray ionization mass spectrometry (ESI-MS) and [13C2]glycine as the donator of two labeled carbon atoms to a purine ring accomplished this goal. We applied the method to quantify purine concentration and *de novo* purine synthesis in placental explants and epithelial adenocarcinoma MCF-7 cells cultivated in the presence of homocysteine. The concentration simulated that at mild and intermediate hyperhomocysteinemia typical for pregnancy-associated complications affecting human placenta, and for breast cancer [14–16]. The multiple manifestations of phenotypic changes under hyperhomocysteinemia are fully described in [10], but the molecular mechanisms of these changes are mostly unknown.

## Materials and methods

### Reagents and reference materials

Solvents and reagents (HPLC grade acetonitrile, methanol, formic acid, ammonium acetate, potassium hydroxide, and perchloric acid) used for sample preparation and as mobile phases were from RSI Labscan LTD (Bangkok, Thailand) and utilized without further purification. Adenine, Guanine, and Hypoxanthine standards, L-glutamine, sodium bicarbonate, [13C2]glycine (cat. num. 283827), and Minimum Essential Medium Eagle (cat. num. M0275) were from Sigma Aldrich (Taufkirchen, Germany). Heat-inactivated fetal bovine serum (cat. num. 12484) was from Thermo Fisher Scientific (Waltham, MA, USA). Polystyrene tissue culture plates (cat. num. 662102) were from Greiner Bio One Cellstar^®^ Tissue Culture Plates (Waltham, MA, USA).

### Instrumentation

An Agilent 1200 Infinity series HPLC system consisted of a G1312 binary pump, a G1322A vacuum degasser, and a G1316A thermostated column compartment (Agilent Technologies, Santa Clara, CA, USA) in combination with a Leap CTC PAL autosampler (Carrboro, NC, USA). The HPLC system was interfaced with an Agilent 6130 single quadrupole massspectrometer (Agilent Technologies, Santa Clara, CA, USA) operating with an electrospray ionization source (ESI) using nitrogen (purity: 99.99%). The equipment was kindly offered by two analytical centers for mass spectrometry equipment of the National Academy of Sciences of Ukraine, in the Institute of Hydrobiology and in the D.K. Zabolotny Institute of Microbiology and Virology. For the chromatographic separation, we tested with two columns: Zorbax-C18 (3.5 × 150 mm, 3.5 μm) and Zorbax SB-Aq C18 (4.6 ×150 mm, 5.0 μm) (Agilent Technologies, Santa Clara, CA, USA).

### Sample collection

The human placental explants and MCF7 cells were used in the study according to the principles of the Declaration of Helsinki. The ethics committee of the Institute of Molecular Biology and Genetics of the National Academy of Sciences of Ukraine (Kyiv) approved the study protocol and the use of human tissues. Placental tissues from elective termination of physiological pregnancies between 8 – 10 weeks of gestation (n = 6) were collected in Kyiv City Clinical Hospital #2 and full term placental samples at 40 weeks of physiological pregnancies (n = 3) were collected immediately after delivery in Kyiv Maternity Hospital #3. Written informed consent was obtained from all participating women. MCF7 cells were from a collection of cell lines in the Institute of Molecular Biology and Genetics of the National Academy of Sciences of Ukraine (Kyiv).

### Cultivation of explants and MCF7 cells

Individual clumps of chorionic villi from the first and third trimester placental samples (ca. 50 – 100 mg) were dissected in sterile cold PBS under a microscope; transferred to the 1 x MEM media (cat. num. M0275) supplemented with 0.292 g/L L-glutamine, 2.2 g/L sodium bicarbonate, and 10 % dialyzed heat-inactivated fetal bovine serum (FBS) in 24-well plates; and cultivated at 37 °C in a humidified atmosphere of 20 % O2 and 5 % CO2 for 24 h [17]. MCF7 cells were cultivated in 100 x 21 mm disposable Petri dishes under the same conditions as the explants, until the cells formed a confluent monolayer, after nearly 4 days. At the beginning of cultivation, the [13C2]glycine was added to final concentration of 80 μM which fully replenished the glycine lacking in the #MO275 MEM media. FBS was dialyzed against 100-fold volumes of PBS at 4 °C for 2 days using a 25 kDa molecular weight cut-off (MWCO) dialysis membrane with daily exchange of dialysis solution [18]. In the control group, the explants and MCF7 cells were cultivated under conditions noted above, and in the experimental group in the presence of 20 μM and 40 μM homocysteine. After cultivation, the explants were carefully gathered, dried on sterile paper towel, put in Eppendorfs, weighed, and stored at – 20 °C until the main procedure. The adhered MCF7 cells were detached from the dish with a 0.25 % trypsin/EDTA solution for 1 min at 37 °C, counted using a hemocytometer, and subjected to centrifugation at 1500 g for 5 min at 4 °C. The cell pellet was stored at – 20 °C until preprocessing before nucleotide base quantification.

### Preprocessing of samples

Preprocessing of placental explants and pellets of MCF7 cells was conducted according to the protocols of Dr. W. Wang in [19]. The samples were suspended in 0.4 ml of 0.4 M perchloric acid, treated in MSE Soniprep 150 Plus ultrasonic disintegrator (MSE Centrifuges, Heathfield UK) for 1 min at 23 KHz and 150 W, and subjected to hydrolysis in a boiling water bath for 60 min to break the glycosidic bonds between the purine base and ribose/deoxyribose group. After the treatments, the samples were cooled for 5 min and neutralized with 5 M KOH in a water-ice bath. The insoluble pellets were removed by centrifugation at 11,000 g, 4 °C for 10 min. The supernatants were collected and stored at –20 °C until LC/MS assay. To control for variations in the recovery of analytes and of ionization efficiency, 200 ng of caffeine (1,3,7-Trimethylpurine-2,6-dione) as an internal standard was added to each sample before hydrolysis.

### HPLC-MS assay for quantification of newly formed and total purines

The qualitative and quantitative detection of purines was conducted by high-performance liquid chromatography – mass spectrometry with electrospray ionization (HPLC/ESI/MS) using single quadrupole mass-spectrometers and UV-Visdiode array detectors (DAD). The hydrolysates of the samples (50 μl) were injected into a rapid resolution **C18 Zorbax column,** which was eluted over 10 min at a flow rate of 0.5 ml/min, with thermostat temperature –30 °C and DAD signal detection at 254 – 260 nm, using an isocratic mobile phase containing 60% of solution A (0.1 M NH_4_Ac with 10% methanol brought to pH 5.0 with formic acid) and 40% of solution B (deionized water).

The **Zorbax SB-Aq C18 column** was injected with the samples (6 μl) and was eluted by a gradient mobile phase consisting of solution A (0.1 M NH_4_Ac with 10 % methanol brought to pH 5.0 with formic acid) and solution B (100 % Methanol). At 0 min the mobile phase contained 80 % A & 20 % B; at 6 min – 80 % A & 20 % B; at 12 min and at 25 min – 100 % solution A. The flow rate was 0.6 ml/min, with thermostat temperature – 32.5 °C. The pH value of eluents was chosen experimentally for optimal balance between detection sensitivity and separation capacity of the columns.

The column eluates were injected directly into a mass spectrometer, which was operated in SIM mode (selected ion monitoring). The positive electrospray mode was used to detect protonated molecular ions [M+H]^+^ of adenine, guanine, and hypoxanthine; negative mode was used to detect deprotonated ion [M-H]^-^ of caffeine in a separate run.

The analytes were quantified using the calibration curves based on mean values of triplicate measurements for five different concentrations of adenine, guanine, and hypoxanthine over the range of 1 – 300 μmol/L by plotting the peak areas against the amount of each injected standard. The linearity of the method was verified by calculation of the regression line using the least squares method.

Limits of detection (LOD) were found by injection of standard solutions of analytes at decreasing concentrations, until the signal to noise ratio (S/N) fell to below 3:1 for an individual analyte. Limits of quantification (LOQ) were calculated as 10 times the signal of detection.

The stock solutions of standards (adenine, guanine·HCl, hypoxanthine, and caffeine) were prepared by dissolving each base in concentrated formic acid (200 μl/mg of base), diluted to concentration 1 mg/ml by the buffer – 0.1 M NH_4_Ac, 10 % methanol, adjusted to pH 5.0 by formic acid, and neutralized with 5 M KOH to pH 5.0. The final concentration 50 μg/ml was adjusted with NH_4_Ac buffer (0.1 M NH_4_Ac, 10 % methanol, pH 5.0). The stock solutions were stored at +4 °C. (N.b. storage at a temperature below +4 °C might induce precipitation of analytes).

### Assay validation

The assay was validated following the FDA guidelines on Bioanalytical Method Validation [20]. Compound stability was assessed by performing several runs for standard adenine, hypoxanthine, and guanine at concentration 40 μmol/L each during one month of stock standard storage at +4 °C.

The intraday and interday variations of this method were measured by performing triple runs of pure analytes and the preprocessed biological samples on the same day and on different days within one month, respectively. Carryover was assessed by running a sequence of a blank sample (deionized water), standard analytes, and a blank sample again. The variations in recovery were controlled by 200 ng of caffeine (1,3,7-Trimethylpurine-2,6-dione, 195 m/z) as an internal standard, which was added to each sample before hydrolysis.

### Mass spectrometry data analysis

The results of the HPLC/ESI-MS assay were processed with the LC/MSD ChemStation software package (Agilent Technologies, Santa Clara, California, USA). The concentration of purines in placental explants was presented in μmoles per 1g of wet tissue. To make the obtained results from both types of biological samples comparable, we presented the purines amount in MCF7 cells per 10^8^ cells as this value corresponds to 1 g of epithelial tissue [21].

### Statistical analysis

All data were tested for normality with Shapiro-Wilk’s W test; they showed a normal distribution (p > 0.05) and were analyzed with statistical parametric methods (see Supplement). The statistical significance of the difference between the levels of purines in explants from the first trimester of gestation cultivated with and without homocysteine was assessed with a one-way ANOVA and a *post hoc* Newman-Keuls test a multiple comparison procedure for the special case of exactly three groups [22]. The statistical significance of the difference between the level of purine bases in MCF cells cultivated with and without homocysteine was assessed by a dependent T-test. The difference between the values in placental explants from the first and third trimesters of gestation cultivated without homocysteine was asseseed by a T-independent test. All the data were represented by mean values with standard errors. Statistical analysis was performed with STATISTICA 10 Enterprise 10.0.1011.6, StatSoft, Inc.

## Results

### Assay establishment and validation

In this study, a simplified HPLC/ESI-MS procedure was developed, established, and applied for the quantification of total and *de novo* synthesized adenine, guanine, and hypoxanthine in placental explants and MCF7 cells. There were two challenges in establishing the assay: a) to find an optimal pH for the mobile phase providing an optimal balance between ESI efficiency of analytes and column separation capacity; b) to enable the accurate simultaneous detection of trace concentrations of labeled purines and the amount of unlabeled purines. To get the optimal separation between protonated ions of adenine (m/z = 136) and hypoxanthine (m/z = 137), we started from a mobile phase with pH 2 and gradually raised pH until pH 5, when sufficient separation was achieved. The second challenge was solved during construction of standard curves (see the next section).

Chromatograms of hypoxanthine, guanine, and adenine mixes were obtained with Zorbax C18 and Zorbax SB-Aq C18 columns and single quadrupole MS systems in SIM mode. The corresponding retention times (RT) of analytes were: 8.542 & 12.113 min for adenine (136 m/z), 4.520 & 5.016 min for guanine (152 m/z), and 4.450 & 4.595 min for hypoxanthine (137 m/z) from Zorbax C18 and Zorbax SB-Aq C18 columns, respectively (Figure 1). The minor peak of adenine with RT = 8.561 min in fig. 1 represented natural M+1 isotope (137 m/z). Both columns very clearly separated adenine from hypoxanthine and from guanine.

**Figure 1.**
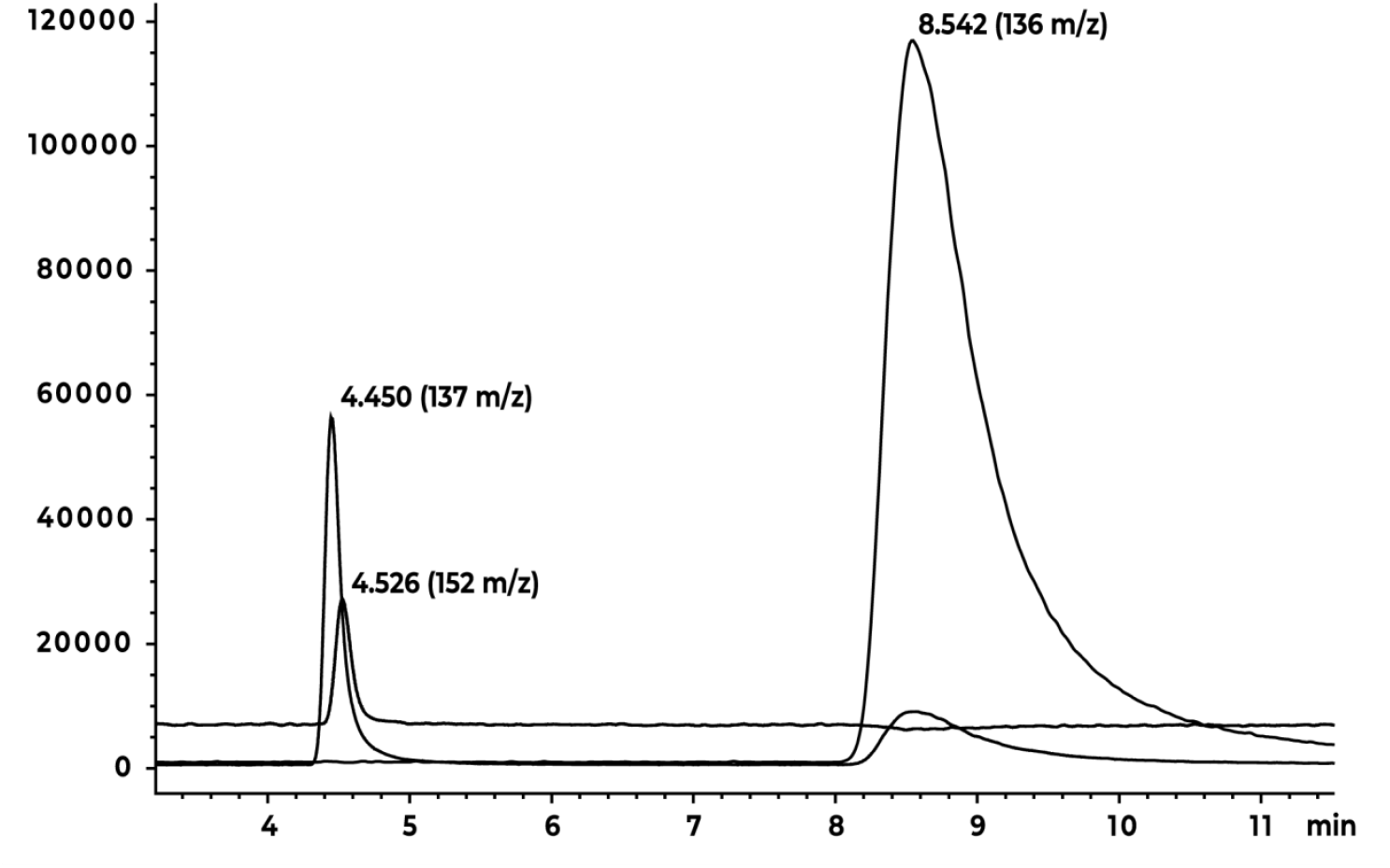
Chromatogram of hypoxanthine (RT 4.450 min, 137 m/z), guanine (RT 4.526 min, 152 m/z), and adenine (RT 8.542 min, 136 m/z) standards (100 ng each nucleotide base in injection) obtained at Zorbax C18 column.

However, the separation of hypoxanthine (m/z = 137) and guanine (m/z = 152) was less effective despite the great difference in their m/z values. We suggest that the similarity in the structure of both compounds rendered their separation difficult (Figure 2).

**Figure 2.**
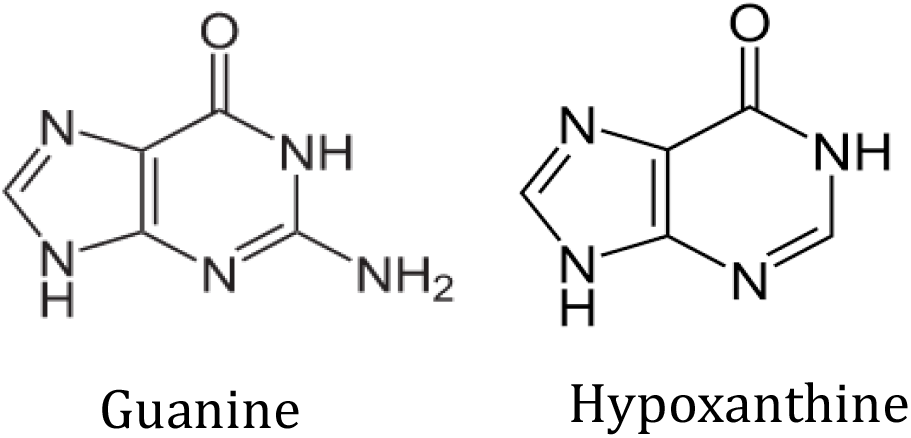
The structure of guanine and hypoxanthine

The separation capacities of the Zorbax C18 and Zorbax SB-Aq C18 columns were principally the same and we chose the Zorbax C18 column for further measurements.

The tests for assay validation showed that signal intensities of standard solutions of adenine, hypoxanthine, and guanine did not change during one-month storage at +4 °C. The intraday repeatability of the HPLC–MS assay was estimated by three consecutive runs of the mixture of standards, 40 μmol/L of each compound. The obtained CV values were 2.9 % for adenine (40 ± 1.2 μmol/L), 3.2 % for hypoxanthine (40 ± 1.3μmol/L), and 4.0 % for guanine (40 ± 1.6 μmol/L). The interday repeatability of the HPLC/ESI-MS assay was demonstrated on three different days within a month using the same concentrations of analytes as for the intraday test. The obtained CV values were higher than for intraday tests: 4.5 % for adenine (40 ± 1.8 μmol/L), 5.0 % for hypoxanthine (40 ± 2.0 μmol/L), and 6.0 % for guanine (40 ± 2.4 μmol/L). The analytes recovery in biological samples comprised 96 – 98 %, estimated according to caffeine recovery as an internal standard with a variability of recovery at CV = 2.43 %. There was no detectable carryover in deionized water according to standards or biological samples.

### Performance characteristics of HPLC/ESI-MS assay for purines quantification

The standard curves were determined for quantification of purines. The linearity of curves was verified by calculation of the regression line using the least squares method. Because of a wide range of concentration intervals and the decrease of MS sensitivity with the increasing amount of injected analyte, we obtained a non-linear MS response to serial dilutions of each individual compound at the highest concentrations. We fit a second-degree polynomial to the data, as is frequently used for such MS analyses [23, 24]. The method showed a good fit (Table 1).

**Table 1.**
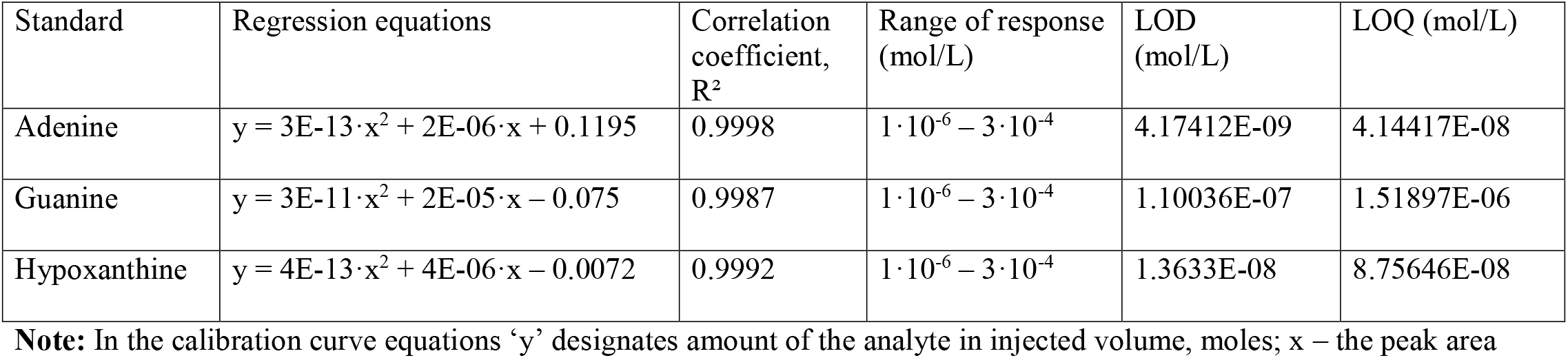
Performance characteristics of purines quantification (Zorbax C18 column)

### Concentration of labeled and unlabeled purines in placental explants and MCF7 cells

The chromatograms of adenine, guanine, and hypoxanthine in both types of biological samples revealed the same retention time as analytes in standard solutions. The mass-detector reliably distnguished heavy-labeled (+2 m/z) adenine (138 m/z), hypoxanthine (139 m/z), and guanine (154 m/z) from the corresponding unlabeled nucleotide bases with 136, 137, and 152 m/z (Figure 3).

**Figure 3.**
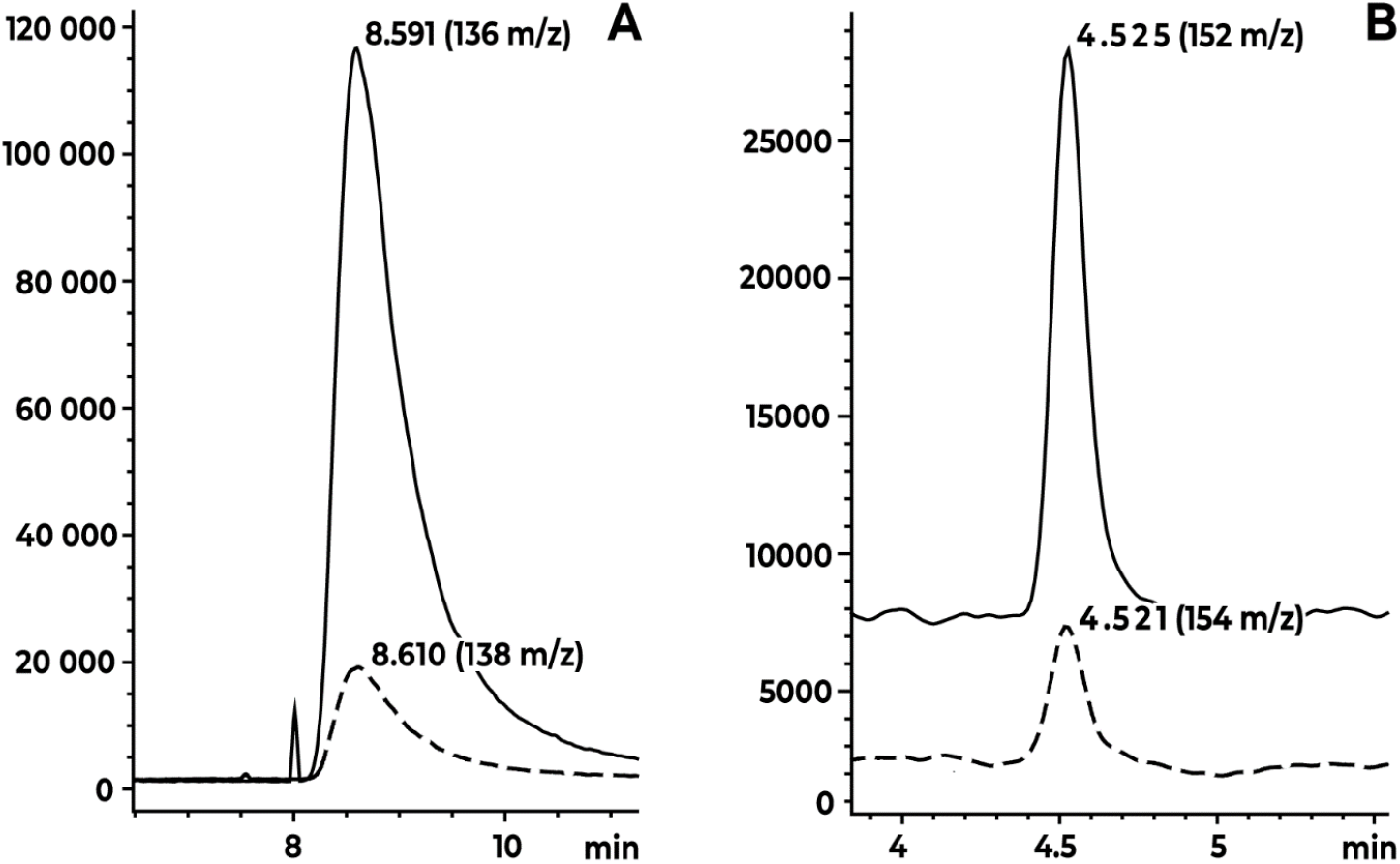
Unlabeled (solid lines) and labeled (dashed lines) adenine (A) and guanine (B) in hydrolysates of placental explants cultivated without homocysteine

The unlabeled adenine, guanine, and hypoxanthine have their natural isotopes with specific m/z values. To calculate the relative abundance of natural isotopes M+2 of each analyte, we used the online tool “Isotope Distribution Calculator and Mass Spec Plotter” (Scientific Instrument Services by Adaptas Solutions, Palmer, MA, USA) available at https://www.sisweb.com/mstools/isotope.htm). For adenine, guanine, and hypoxanthine the M+2 / M ratios comprise 0.233, 0.437, and 0.411 %%, respectively. As peaks of natural isotopes M+2 overlap with the peaks of [13C2]labeled analytes, concentrations of the latter were calculated taking into account the estimated peak areas of natural isotopes M+2.

The quantification of labeled and unlabeled purines revealed that all obtained values were above LOQ values, except labeled hypoxanthine and labeled guanine in several samples. The concentration of unlabeled purines in placental explants from the first and third trimesters of gestation and in MCF7 cells, either treated and not treated with homocysteine, were of the same order of magnitude; arranged in descending order of the values: adenine > guanine > hypoxanthine. Despite a similar concentration of unlabeled purines, the concentrations of newly synthesized (labeled) adenine, guanine, and hypoxanthine and their combined fractions were more than one order of magnitude less in placental explants than in MCF7 cells (Tables 2, 3, Supplememt).

**Table 2.** Concentration of labeled and unlabeled purine bases in placental explants from first trimester of gestation (μmoles/g tissue)

| Hcy.<br>$\mu\text{M}$ | A | A** | Hyp | Hyp** | G | G** | Labeled<br>purines | Total<br>purines | Labeled<br>pur/<br>Total pur.,<br>%% |
| --- | --- | --- | --- | --- | --- | --- | --- | --- | --- |
| 0 | 4.71<br>$\pm 0.283$ | 0.05<br>$\pm 0.006$ | 0.028<br>$\pm 0.001$ | n.d. | 4.0<br>$\pm 0.32$ | 0.04<br>$\pm 0.001$ | 0.10<br>$\pm 0.006$ | 8.8<br>$\pm 0.50$ | 1.06<br>$\pm 0.132^{\gamma}$ |
| 20 | 3.8<br>$\pm 0.27$ | 0.03<br>$\pm 0.002^*$ | 0.021<br>$\pm 0.002$ | n.d. | 2.8<br>$\pm 0.21$ | 0.02<br>$\pm 0.004$ | 0.05<br>$\pm 0.005$ | 6.7<br>$\pm 0.47$ | 0.75<br>$\pm 0.155^{\gamma} *$ |
| 40 | 2.9<br>$\pm 0.14$ | 0.02<br>$\pm 0.001^*$ | 0.018<br>$\pm 0.002$ | n.d. | 1.5<br>$\pm 0.09$ | 0.001<br>$\pm 0.0001$ | 0.022<br>$\pm 0.001$ | 4.4<br>$\pm 0.24$ | 0.49<br>$\pm 0.008^*$ |

**Table 3.**
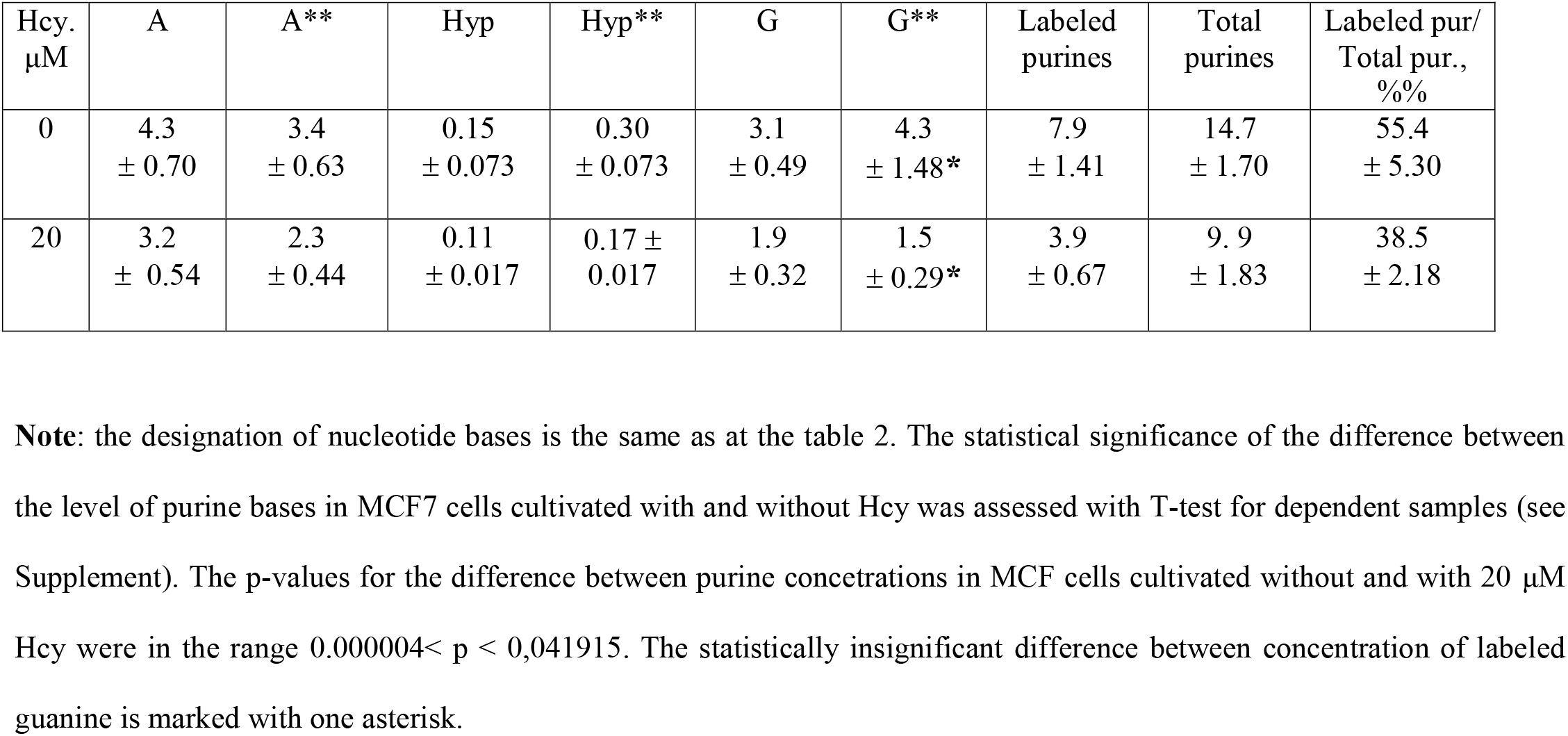
Concentration of labeled and unlabeled purine bases in MCF7 cells (μmoles/10^8^ cells)

The cultivation of both biological samples with 20 μM homocysteine caused an approximately two-fold decrease in the concentration of total newly synthesized purines and ~1.5 fold decreases in the concentration of total (labeled plus unlabeled) purines. The cultivation of placental explants with 40 μM homocysteine induced more pronounced changes in the concentration of unlabeled and labeled purines than cultivation with 20 μM homocysteine (Tables 2, 3, Supplement).

We have analyzed how purine biosynthesis changes in human placenta at the end of gestation in comparison with the first trimester. The concentration of all types of purines in placental explants from the third trimester of gestation (Table 4) was substantially less than in the first (Table 2, first line), and nearly equal to that in placental explants from the first trimester cultivated with 40 μM homocysteine (Table 2, third line). The percentage of newly formed purines among total purines was 0.55 ± 0.004 % in explants from the third trimester, and 0.49 ± 0.008 % in the first trimester explants cultivated with 40 μM homocysteine

**Table 4.**
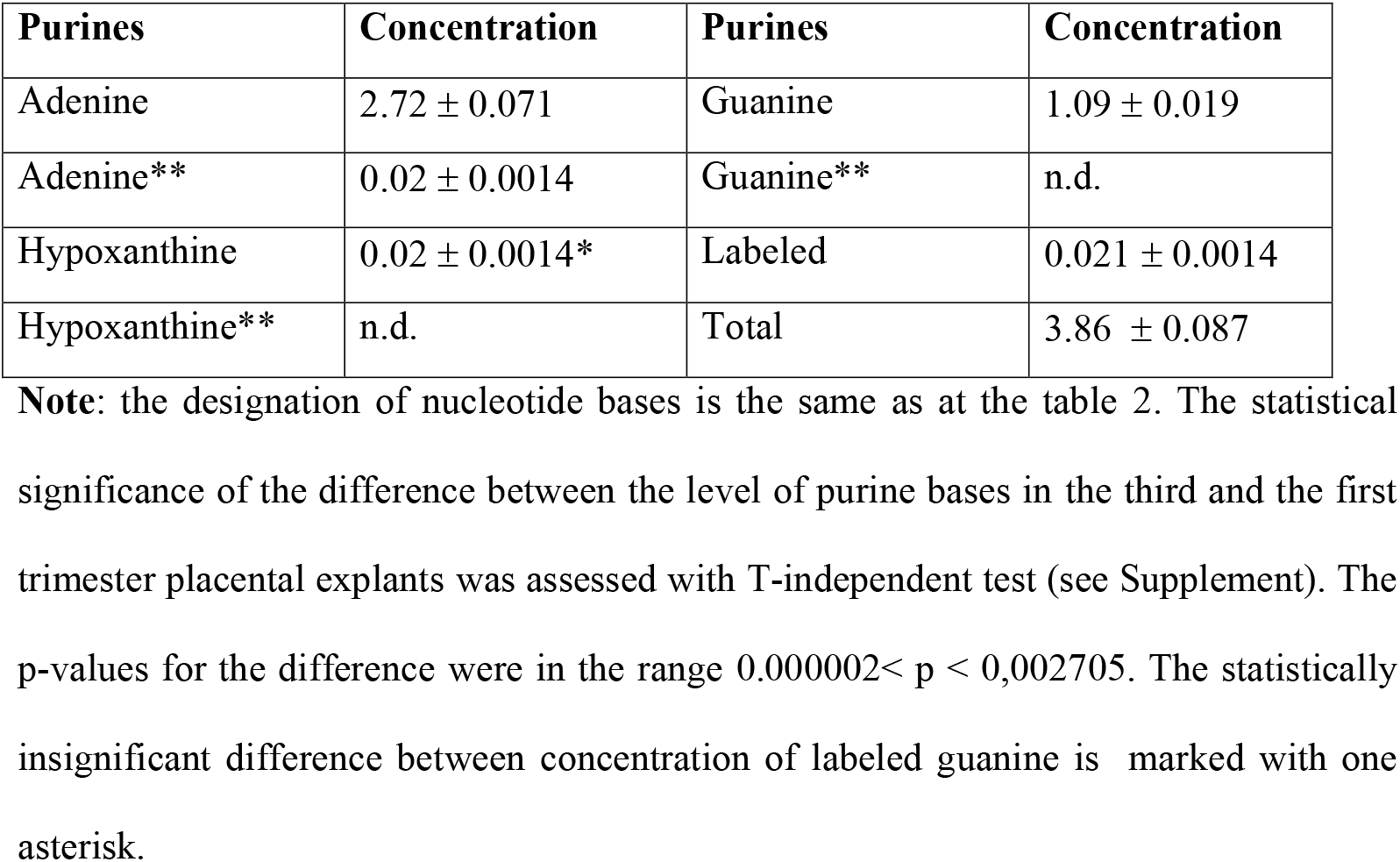
Concentration of labeled and unlabeled purine bases in placental explants from the third trimester of gestation (μmoles/g tissue)

## Discussion

We have established and tested a simplified method for the quantification of total and newly synthesized purines, and applied it to corresponding measurements in human placental explants cultivated with homocysteine, which imitated mild and intermediate hyperhomocysteinemia typical for preeclampsia. Epithelial adenocarcinoma MCF7 cells were used for comparison. The total (labeled and unlabeled) purine concentration reflected the total cellular/tissue concentration of nucleotide bases, and included free acid soluble bases and the bases released by acid hydrolysis from free nucleosides, nucleotides, and their derivatives, and from DNA, RNA, and oligonucleotides. The amount of newly labeled purines in relation to the total purine content reflected the rate of *de novo* purine synthesis.

### Simplified HPLC/ESI-MS method vs. similar approaches

This simplified method of purine quantification is robust, fast, easy, safe, and avoids tedious steps or costly reagents; it is applicable to measurement in a wide range of concentrations. The hand-on processes take around 1.5 hr. The whole procedure requires only customary equipment for a biochemical laboratory plus an LC-MS analysis instrument which is substantially cheaper than an LC-MS/MS. Stable [13C2]glycine, in contrast to radioactive glycine, is fully compatible with live cells, and may completely replace unlabeled glycine. The trophoblast and MCF7 cells take up glycine [7, 25], which contributes adjacent C-4, C-5, and N-7 atoms to a purine base ring, and in the case of stable [13C2]glycine makes a purine ring heavier by +2 m/z units and detectable by MS.

In experiments addressing similar questions, researchers frequently used radioactive labels and presented results in cpm/mln of cells, which are not easily comparable to other measurements [12, 19, 26]. Only two publications took into account all cellular purine nucleotides including acid-soluble and non-acid-soluble pools [19], while another presents the data in fractions of cellular nucleotides, which do not reflect the total rate of purine synthesis [27, 28]. The overwhelming majority of studies used cultivated cells [12, 19, 26]. Data on purine content and synthesis in human tissues, or tissues of laboratory animals not affected by tumors, are simply absent.

### Rate of purine synthesis in placental explants: the impact of homocysteine and gestational age

This analysis of purine concentration and the rate of purine synthesis in placental explants and MCF7 cells has revealed great difference between both types of samples, especially between their fractions of newly synthesized purines. MCF7 human mammary carcinoma cells, after a lag period of about 48 hr., grow exponentially during 4 – 5 days with a mean population doubling time of about 24 hr., and 80 to 90 % of cells in the rapidly cycling pool until formation of a monolayer [29]. Therefore, the 50 % fraction of newly synthesized purines in MCF7 cells after 4 days of growth was fully expected. The situation in placental explants was substantially different. A single layer of dividing cytotrophoblast that lines the stromal cell core differentiates into syncytiotrophoblast, a non-dividing layer, and extravillous cytotrophoblast, which forms trophoblast cell columns and invades the decidual interstitium along the endovascular, endolymphatic, and endoglandular routes [30]. As a result, the fraction of proliferating cells in placental explants is significantly restricted compared to MCF7 cells.

The total purines and the fraction of newly synthesized purines distinctly decreased in full-term explants compared to first trimester explants. These data supplemented our previous studies which revealed the lower abundance of *GART* and *ATIC* mRNAs encoding enzymes involved in the synthesis of a purine ring [31]. The decrease of *de novo* purine synthesis corroborated the decreased intensity of DNA synthesis in fullterm placenta, which is demonstrated by proliferation markers Ki67 and MIB-1 [32, 33]. All the traits together make a substantial contribution to the general down-regulation of proliferating activity as the gestational age increases. It is noteworthy that the concentration of total and newly synthesized purines in full-term placenta is similar to the concentration of corresponding purines in first trimester explants cultivated with 40 μM homocysteine.

The mild or intermediate hyperhomocysteinemia is characteristic of many human diseases, including breast cancer and preeclampsia [14–16]. Cultivation of both types of samples in the presence of homocysteine in the cultivation medium, which imitated the mild or intermediate hyperhomocysteinemia, resulted in the down regulation of the *de novo* purine synthesis. This effect of homocysteine in placental explants is consistent with our prediction of a 30% restriction of metabolic fluxes through purine *de novo* biosynthesis under double the homocysteine load of a placental stoichiometric model of folate metabolism [34], and with the decrease of Ki67 marker abundance in placental explants under the same homocysteine treatment [35]. Therefore, the decline of purine synthesis corroborates the general down-regulation of proliferating activity in response to homocysteine.

Purine *de novo* biosynthesis is an integral part of the folate cycle. Homocysteine is an intermediate metabolite in the methionine cycle. Both cycles refer to folate-mediated one carbon metabolism and are tightly bound via the reaction of homocysteine remethylation to methionine. In placenta, methyltetrahydrofolate from the folate cycle (and not betaine!) serves as a sole methyl group donor to homocysteine to form methionine, while released tetrahydrofolate is used in the folate cycle and supports its reactions, including purine and pyrimidine biosynthesis. In the methionine cycle, methionine is subjected to sequential reactions with the following intermediate products: methionine → S-adenosylmethionine (SAM) → S-adenosylhomocysteine (SAH) and, finally, SAH hydrolysis to homocysteine and adenosine. In the majority of tissues, about 50% of homocysteine is conserved by remethylation to methionine, and nearly the same amount is irreversibly converted by transsulfuration pathway to cysteine, the precursor of glutathione, taurine, and hydrogen sulfide along the pathway of cysteine catabolism [36, 37]. Due to the interaction of the folate and methionine cycles, we discuss the metabolic fluxes through these cycles to understand the mechanism of homocysteine’s impact on *de novo* purine biosynthesis.

Our previous studies revealed that the double homocysteine load in the placental stoichiometric model of folate-mediated one carbon metabolism induced a 12 % elevation of metabolic flux throughout the methionine cycle [34]. In our previous experiments, the cultivation of placental explants with 20 μM homocysteine nearly doubled the concentration of SAM and SAH with an unchanged SAM/SAH ratio [38], activated cystathionine β-synthase (EC 4.2.1.22), and increased production of cysteine as compared to explants cultivated without homocysteine [39]. Cystathionine β-synthase is a heme-based redox sensor, which modulates its activity in response to changes in the ambient redox potential. Cystathionine β-synthase has a remarkably high Km for homocysteine that is in the millimolar range [40]. This is why cystathionine β-synthase is sensitive to elevated concentrations of homocysteine and actively converts it to cysteine, the limiting reagent in the synthesis of glutathione that would correct an oxidative disturbance induced by homocysteine. These experimental results point to the intensified metabolism of homocysteine in the methionine cycle, and thereafter homocysteine clearance by transsulfuration under a homocysteine load. The folate-utilizing enzymes actively compete for folate derivatives as enzymatic binding capacity is far in excess of the restricted amount of folate derivatives [41]. Cytoplasmic C-1-tetrahydrofolate synthase (MTHFD1) is a trifunctional enzyme responsible for formation of 5,10-methylenetetrahydrofolate, 5,10-metenyltetrahydrofolate and formyltetrahydofolate. The first derivative from this list is reduced to 5-methyltetrahydofolate, which provides a methyl group to homocysteine for methionine synthesis while the last one is the donor of formate to purine biosynthesis. 5,10-metenyltetrahydrofolate is an intermediate between the former and the latter folate derivatives during their reversible interconversion [42]. We suggested that the elevated SAM and SAH concentration in placental explants under 20 μM homocysteine treatment argues for preferential formation of 5,10-methylenetetrahydrofolate and its usage for reduction to 5-methyltetrahydofolate and further activation of methionine cycle at expense of other MTHFD1 activity supporting the synthesis of formyltetrahydofolate required for *de novo* purine synthesis.

According to our knowledge, it is the first experimentally and bioinformatically based discussion of homocysteine impact on the shift of metabolic fluxes in placental explants. The impact of homocysteine results in down regulation of *de novo* purine biosynthesis and might have adverse consequence for placental growth and development.

The changes in placental explants and MCF7 cells induced by homocysteine partly resemble down-regulation of *de novo* purine synthesis in vascular endothelial cells, hepatocytes and neuronal precursors under the influence of hyperhomocysteinemia [10] and principally differ from enhanced proliferation of vascular smooth muscle cells under the same treatment [11]. The additional investigations are required to elucidate the mechanism of tissue-specific activity of homocysteine regarding *de novo* purine biosynthesis.

## Limitations of the assay

We characterized the concentration of total and newly formed purines in placental explants and MCF7 cells and did not characterize the salvage pathway though *de novo* and salvage pathways make their input in concentration of total purines.

The analytes were measured in ESI+ and the caffeine IS was measured in ESI-and in a separate run. It would have been better to use an IS that can be detected simultaneously with the analytes as it could have been corrected much better for any sample preparation variations.

The usage of HPLC/ESI-MS for measurement the concentration of total and newly synthesized purines has several advantages mentioned in the text. The usage of HPLC/ESI-MS/MS would provide the increased sensitivity and better separation of hypoxanthine and guanine

## Conclusions

This work represents a step forward in the effort to improve, standardize, and increase the adoption of high performance liquid chromatography associated with mass spectrometry and electrospray ionization for quantification of total and *de novo* synthesized purines in human tissues especially in low mg weight. Usage of stable isotope dramatically improves the convenience, reliability, and ease of use of the analysis. These advantages provide the opportunity for more labs to explore the tissue specific aspects of purine biosynthesis even at low activity. In view of the search of new tissue specific targets for anticancer therapy, the most challenging aspect of purine synthesis is its specific regulation in actively proliferating different tissues.

The next major advancement of this study is the demonstration of metabolic shift in folate – mediated metabolism in placental explants under homocysteine treatment, which imitates mild and intermediate hyperhomocysteinemia. The shift to methionine synthesis and transsulfuration at expense of purine synthesis is suggested to be one of the mechanisms of adverse effect of hyperhomocysteinemia on placental development and function.

## Acknowledgements

We are grateful to Mr. Kenneth Sheridan and to Dr. Pavel Shliaha (Sloan Kettering Institute, NY, US) for critical reading of the manuscript and helpful comments. We would like to thank the patients and clinicians without whom this study would not have been possible. The study was supported by the National Academy of Sciences of Ukraine (Project 0115U003748).

## Compliance with Ethical Standards

### Conflict of Interest

The authors declare that they have no conflicts of interest. The human placental explants cells were used in the study according to the requirements of the Declaration of Helsinki, 1964. The ethics committee of the Institute of Molecular Biology and Genetics of the National Academy of Sciences of Ukraine (Kyiv) approved the study protocol and the use of human tissues. Placental tissues were collected in Kyiv City Clinical Hospital #2 and Kyiv Maternity Hospital #3; written informed consent was obtained from all participating women.

## Author Contributions

RR and SR conceived and designed research. IK and MK conducted MS analysis, VK conducted MCF7 cells cultivation, IV provided samples, RR, SR, and AK conducted wet experiments, MO discussed results, MO and RR wrote the manuscript. All authors read, discussed, and approved the manuscript.

## Notes

### Competing Interest Statement

The authors have declared no competing interest.

## References

1. Nelson D.L., Cox M.M.: Lehninger Principles of Biochemistry. Chapter 8. Nucleotides and Nucleic Acids. 7th edition. 1271 pp. W.H. Freeman & Co. Ltd. (2017)

2. Lane, A. N., Fan, T. W.-M.: Regulation of mammalian nucleotide metabolism and biosynthesis. Nucleic Acids Res. 43(4), 2466–2485 (2015)

3. Burnstock, G., Boeynaems, J.-M.: Purinergic signaling and immune cells. Purinergic Signal. 10(4), 529–564 (2014)

4. McCarthy, M. T., Moncayo, G., Hiron, T. K., Jakobsen, N. A., Valli, A., Soga, T., O'Callaghan, C. A.: Purine nucleotide metabolism regulates expression of the human immune ligand MICA. J. Biol, Chem. 293(11), 3913–3924 (2018)

5. Pedley, A. M., Benkovic, S. J.: A New view into the regulation of purine metabolism: the Purinosome. Trends Biochem. Sci. 42(2), 141–154 (2017)

6. Momb, J., Lewandowski, J. P., Bryant, J. D., Fitch, R., Surman, D. R., Vokes, S. A., Appling, D. R.: Deletion of Mthfd1l causes embryonic lethality and neural tube and craniofacial defects in mice. Proc. Natl. Acad. Sci. U S A. 110(2), 549–54 (2013)

7. Kovalchuk, V., Samluk, Ł., Juraszek, B., Jurkiewicz-Trząska, D., Sucic, S., Freissmuth, M., Nałęcz, K. A.: Trafficking of the amino acid transporter B0+ (SLC6A14) to the plasma membrane involves an exclusive interaction with SEC24C for its exit from the endoplasmic reticulum. Biochim, Biophys, Acta Mol, Cell, Res, 1866(2), 252–263 (2019)

8. Ben-Sahra, I., Hoxhaj, G., Ricoult, S. J. H., Asara, J. M., & Manning, B. D.: mTORC1 induces purine synthesis through control of the mitochondrial tetrahydrofolate cycle. Science 351(6274), 728–733 (2016)

9. Emmanuel, N., Ragunathan, S., Shan, Q., Wang, F., Giannakou, A., Huser, N., Jin, G., Myers, J., Abraham, R.T., Unsal-Kacmaz, K.: Purine Nucleotide Availability Regulates mTORC1 Activity through the Rheb GTPase. Cell Rep. 19(13), 2665–2680 (2017)

10. Esse, R., Barroso, M., Tavares de Almeida, I., Castro, R.: The Contribution of Homocysteine Metabolism Disruption to Endothelial Dysfunction: State-of-the-Art. Int. J. Mol. Sci. 20(4), 867 (2019)

11. Zou, T., Yang, W., Hou, Z., Yang, J.: Homocysteine enhances cell proliferation in vascular smooth muscle cells: role of p38 MAPK and p47phox. Acta Biochim. Biophys. Sin. 42(12), 908–915 (2010)

12. An, S., Deng, Y., Tomsho, J. W., Kyoung, M., Benkovic, S. J.: Microtubule-assisted mechanism for functional metabolic macromolecular complex formation. Proc. Natl. Acad. Sci. U S A. 107(29), 12872–12876 (2010)

13. Cordell, R. L., Hill, S. J., Ortori, C. A., Barrett, D. A.: Quantitative profiling of nucleotides and related phosphate-containing metabolites in cultured mammalian cells by liquid chromatography tandem electrospray mass spectrometry. J. Chromatogr. B Analyt. Technol. Biomed. Life Sci. 871(1), 115–124 (2008)

14. Hannibal, L., Blom, H. J.: Homocysteine and disease: Causal associations or epiphenomenons? Mol. Aspects Med. 53, 36–42 (2017)

15. Mansour, A., Harb, H., Abdelhafee, M.: Diagnostic Value of Homocysteine and Other Preeclampsia Markers: Relationship with Severity. Int. J. Biol. Chem. 5(4), 227–237 (2011)

16. Naushad, S. M., Reddy, C. A., Kumaraswami, K., Divyya, S., Kotamraju, S., Gottumukkala, S.R., Digumarti, R.R., Kutala, V. K.: Impact of Hyperhomocysteinemia on Breast Cancer Initiation and Progression: Epigenetic Perspective. Cell Biochem. Biophys. 68(2), 397–406 (2014)

17. Baczyk, D., Dunk, C., Huppertz, B., Maxwell, C., Reister, F., Giannoulias, D., Kingdom, J. C. P.: Bi-potential behaviour of cytotrophoblasts in first trimester chorionic villi. Placenta. 27(4-5), 367–374 (2006)

18. An, S., Kumar, R., Sheets, E., Benkovic, S.: Reversible compartmentalization of de Novo Purine Biosynthetic Complexes in Living Cells. Science 320(5872), 103–106 (2008)

19. Wang, W., Fridman, A., Blackledge, W., Connely, S., Wilson, I., Pilz, R., Boss, G.: The phosphatidylinositol 3-kinase/Akt cassette regulates purine nucleotide synthesis. J. Biol, Chem. 284(6), 3521–3528 (2009)

20. Zimmer, D.: New US FDA draft guidance on bioanalytical method validation versus current FDA and EMA guidelines: chromatographic methods and ISR. Bioanalysis. 6(1), 13–19 (2014)

21. Del Monte, U.: Does the cell number 10(9) still really fit one gram of tumor tissue? Cell Cycle 8(3), 505–506 (2009)

22. Seaman, M. A., Levin, J. R. & Serlin, R. C. New Developments in pairwise multiple comparisons: Some powerful and practicable procedures Psychol. Bull. 110(3), 577–586 (1991).doi: 10.1037/0033-2909.110.3.577

23. Hinshaw, J. V.: Nonlinear calibration. LCGC. 20(4), 120–126 (2002)

24. Kirkup, L., Mulholland, M.: Comparison of linear and non-linear equations for univariate calibration. J. Chromatogr A. 1029(1-2), 1–11 (2004)

25. Holm, M. B., Bastani, N. E., Holme, A. M., Zucknick, M., Jansson, T., Refsum, H., Michelsen, T. M.: Uptake and release of amino acids in the fetal-placental unit in human pregnancies. PLoS One. 12(10), e0185760 (2017)

26. Fridman, A., Saha, A., Chan, A., Casteel, D., Pilz, R., Boss, G.: Cell cycle regulation of purine synthesis by phosphoribosyl pyrophosphate and inorganic phosphate. Biochem. J. 454(1), 91–99 (2013)

27. Labuschagne, C. F., van den Broek, N. J. F., Mackay, G. M., Vousden, K. H., Maddocks, O. D. K.: Serine, but not glycine, supports one-carbon metabolism and proliferation of cancer cells. Cell Rep. 7(4), 1248–1258 (2014)

28. Zhao, H., Chiaro, C. R., Zhang, L., Smith, P. B., Chan, C. Y., Pedley, A. M., Benkovic, S. J.: Quantitative Analysis of Purine Nucleotides Indicates That Purinosomes Increase de Novo Purine Biosynthesis. J. Biol. Chem. 290(11), 6705–6713 (2015)

29. Sutherland, R. Hall, I. Taylor, Cell proliferation kinetics of MCF-7 human mammary carcinoma cells in culture and effects of tamoxifen on exponentially growing and plateau-phase cells. Cancer Res. 43(9), 3998–4006 (1983)

30. Moser, G., Windsperger, K., Pollheimer, J., de Sousa Lopes, S. C., Huppertz, B.: Human trophoblast invasion: new and unexpected routes and functions. Histochem. Cell Biol. 150(4), 361–370 (2018)

31. Korneyeva, K. L., Rodriges, R. R., Ralchenko, S. V., Vakulenko, A. V., Manzhula, L. V., Melnik, V. T., Vereshchak, O.Y.,. Obolenskaya, M. Y.: The genes expression, which is encoding enzyme key reactions of folate-dependent metabolism in human placenta in the first and third trimesters of uncomplicated pregnancy. Perinatol. Pediatr. 60(4), 24–30 (2014)

32. Martsenyuk, O. P., Mislanova, C., Romanets(Korneeva), K. L., Teplyuk, N. M., Obolenskaya, M. Y.: The level of low molecular thiols and folate in human. Ukr.Biochem. J. 81(4), 94–104 (2009)

33. Smith, S. C., Baker, P. N., Symonds, E. M.: Increased placental apoptosis in intrauterine growth restriction. Am. J. Obstet. Gynecol. 177(6), 1395–1401 (1997)

34. Rodriguez, R. R., Lushchyk, I. S., Obolenskaya, M. Y.: Stoichiometric model of folate-dependent metabolism of one-carbon units in human placenta. Ukr. Biochem. J. 84(4), 20–31 (2012)

35. Martseny uk, O. P., Romanets, K. L. Obolens'ka, M. I., Huppertz, B.: Effect of homocysteine on the structure and functions of human placenta trophoblasts. Ukr. Biochem. J. 81(5), 40–49 (2009)

36. Finkelstein, J. D.: Pathways and regulation of homocysteine metabolism in mammals. Semin. Thromb. Hemost. 26(3), 219–225 (2000)

37. Tarver, H., Schmidt, C. L. A. The conversion of methionine to cystine: experiments with radioactive sulfur. J. Biol. Chem. 130(1), 67–80 (1939)

38. Rodriguez, R., Vakulenko, O., Ralchenko, S., Kostiuk, A., Porublyova, L., Konovets, I., Voronina, I, Obolenskaya, M.: Quantification of S-adenosyl-methionine and S-adenosylhomocysteine in human placenta and placental explants under homocysteine treatment. Intern. J. Mass Spectrom. 421, 279–284 (2017)

39. Mislanova, C., Martsenyuk, O., Huppertz, B., Obolenskaya, M.: Placental markers of folate-related metabolism in preeclampsia. Reproduction, 142(3), 467–476 (2011)

40. Banerjee, R., Zou, C.: Redox regulation and reaction mechanism of human cystathionine-?-synthase: a PLP-dependent hemesensor protein. Arch. Biochem. Biophys. 433(1), 144–156 (2005)

41. Fox, J. T., Stover, P. J.: Folate-mediated one-carbon metabolism. Vitam. Horm. 79(08), 1–44 (2008)

42. Stover, P. J.: One-carbon metabolism-genome interactions in folate-associated pathologies. J. Nutr. 139(12), 2402–2405 (2009)

